# Riverine plastic litter restructures microbial vitamin B_12_ metabolism and generates functional heterogeneity

**DOI:** 10.64898/2026.09.06.749157

**Authors:** Priscilla Carrillo-Barragán, Erik Zettler, Sebastian Rieder, Leslie G. Murphy, Tom Theirlynck, Patricia Burkhardt-Holm, Linda Amaral-Zettler

## Abstract

Rivers transport most land-derived plastic waste to the ocean, yet how this reshapes microbial metabolic potential remains poorly resolved. We characterized plastisphere and water-column communities across 14 stations along the River Rhine, integrating ATR-FTIR polymer characterization, 16S/18S rRNA amplicon profiles, and 120 metagenome-assembled genomes. Plastisphere communities were taxonomically distinct from water communities (PERMANOVA R² = 0.258, p = 0.0001) and, despite no overall shift in functional centroid (R² = 0.245, p = 0.125), were markedly more heterogeneous in functional composition across sites (permutest p = 0.0047). Aerobic corrin ring synthesis, the core B_12_ biosynthetic pathway, was among the most differentially dispersed functions and enriched on plastic at all seven paired stations (Wilcoxon exact test, W = 3, p = 0.004079). This signal coincided with a diatom-dominated eukaryotic plastisphere community; diatoms cannot synthesize B_12_ and depend on bacterial provisioning, linking functional and taxonomic restructuring via a plausible cross-domain mechanism. Plastisphere communities were also less tightly coupled to the river’s dissolved nutrient gradient than water communities (envfit R² = 0.64 vs. 0.82). Together, these results indicate that riverine plastic litter does not merely accumulate biomass passively but actively restructures specific metabolic capacities of its colonizers, exemplified by vitamin B_12_ metabolism.

**Originality-Significance Statement:** This study presents a genome-resolved, transect-scale comparison of plastisphere and free-living microbial communities across a major European river, integrating polymer characterization, rRNA amplicon profiling, and 120 metagenome-assembled genomes. We show that plastic substrates partially decouple community assembly from the river’s dissolved nutrient gradient while selectively enriching vitamin B₁₂ biosynthetic potential, linking this functional restructuring to a co-occurring diatom-dominated eukaryotic community that depends on bacterial B₁₂ provisioning. By identifying a specific, testable metabolic mechanism underlying plastisphere functional distinctiveness, this work moves beyond descriptive taxonomic comparisons toward a mechanistic understanding of how plastic pollution reshapes microbial biogeochemical function in freshwater systems.

## Introduction

Rivers transport approximately 80% of land-based plastic waste to the ocean [1], yet the ecological consequences of this flux for microbial communities colonizing plastic surfaces along the way remain poorly understood. Unlike stationary samples, or samples that are incubated in a set location, large river systems offer continuous gradients of urbanization, nutrient loading, and plastic abundance that stretch hundreds of kilometers, making them ideal systems in which to ask how environmental conditions shape the ecology of plastic-associated biofilms [2–5]. Most existing riverine studies, however, rely on a small number of stations lacking the spatial range or co-sampled environmental data needed to resolve gradient structure, and few couple community-level observations to genome-resolved functional data.

The River Rhine is one of Europe’s most industrialized rivers and an ecologically compelling system in which to address this gap [6, 7]. Flowing through six countries and supporting approximately 50 million inhabitants, the Rhine serves densely urbanized zones across Switzerland, Germany, and the Netherlands. Its basin hosts approximately 10% of the global chemical industry [5], and mean surface microplastic concentrations along the Rhine reach ∼8.9 × 10⁵ particles km⁻² >300 µm [5], with polymer composition dominated by polyethylene (PE), polypropylene (PP), and polystyrene (PS) [5, 7, 8].

Polymer surfaces are hydrophobic, chemically recalcitrant, and enriched with leaching additives and adsorbed contaminants, conditions that initially favor taxa capable of surface attachment, biofilm formation, and xenobiotic metabolism [4, 9–12], though these substrate-specific selective signals often diminish as communities mature [13, 14]. Nonetheless, plastic-associated communities remain detectably distinct from co-occurring free-living communities across aquatic environments [12, 15–17], consistent with the biofilm habitat itself, rather than polymer chemistry alone, imposing selective pressures on community assembly [14]. A key but underexplored consequence is that plastisphere communities may therefore be partially decoupled from the ambient environmental gradients that structure free-living communities and may additionally develop distinct metabolic capacities not captured by taxonomic comparison alone.

One such capacity is the biosynthesis of vitamin B_12_ (cobalamin), an essential cofactor that most eukaryotic algae, including diatoms, cannot synthesize and must instead acquire from co-occurring bacteria [18, 19]. This auxotrophy makes B_12_ provisioning a well-established axis of microbial cross-feeding in aquatic systems [18], and genome-resolved work on marine plastisphere biofilms has shown that transporters and enzymes central to B_12_-dependent metabolism become more abundant as biofilms mature [15]. Two distinct biosynthetic routes to the corrin ring, the biosynthetic core of the B_12_ molecule, exist in bacteria: an aerobic pathway and a phylogenetically distinct anaerobic pathway [20, 21]. Whether plastic substrates in freshwater systems selectively enrich for corrinoid biosynthesis capacity, and whether this maps onto the eukaryotic taxa that depend on it, has not previously been tested along a river continuum.

Riverine plastisphere ecology is a growing field. Pan-European research across nine rivers identified a recurrent plastisphere dominated by Gammaproteobacteria and Bacteroidota, with community composition modulated by nutrient status and geography [22]. In the Dutch portion of the Rhine, microplastics were shown to host distinct bacterial assemblages with reduced diversity compared to surrounding water, enriched in biofilm-forming taxa [23]. Nevertheless, nutrient concentrations, particularly nitrogen and phosphorus, increase along the spatial gradient of the River Rhine which can affect the communities [24]. Direct quantitative comparisons of how plastisphere and free-living communities track longitudinal nutrient gradients along a river main stem, coupled to genome-resolved functional data, remain scarce.

Here we address these questions using plastics collected during the 2019 Greenpeace *Beluga II* expedition, that sampled 14 stations spanning 820 km of the Rhine River main stem between Rotterdam, the Netherlands and Basel, Switzerland. By integrating polymer characterization, rRNA amplicon-based community profiles, and 120 quality-filtered metagenome-assembled genomes, we ask: (1) Do plastic-associated microbial communities show weaker coupling to dissolved nutrient gradients than free-living assemblages?; (2) Does the plastic substrate generate functional heterogeneity beyond what these gradients explain; and (3) Does this heterogeneity converge on a specific metabolic function, corrinoid biosynthesis, that plausibly links the plastisphere’s distinctive taxonomic and functional composition. We interpret our findings in the context of plastic association and functional restructuring in one of Europe’s most anthropogenically influenced river systems.

## Experimental Procedures

### Field sampling and onboard processing

Sampling was conducted once or twice daily at 14 stations between Rotterdam (GP001, hereafter 01) and Basel (GP014, hereafter 14) during the 2019 Greenpeace *Beluga II* Rhine expedition (23 March – 5 April 2019; all station-habitat combinations included in each analysis are listed in Supplementary Table S1). At each station, surface water and plastic debris were collected across multiple size classes. All handling was performed with nitrile gloves.

Surface-water samples were collected during each manta-net tow using acid-rinsed polypropylene buckets. Water subsamples were partitioned for: (i) DNA extraction, filtered through Sterivex cartridges (free-living fraction, ST); and (ii) nutrient analysis (filtered 0.2 µm; NO₃⁻+NO₂⁻, PO₄³⁻, and Si frozen at −20 °C; silicate refrigerated at 4 °C).

Meso- and microplastics were collected with a 2 m outrigger manta net (330 µm mesh) towed for 10–30 min at 1.5–2.5 knots. After retrieval, visible plastic fragments were isolated using ethanol-rinsed forceps, rinsed with 0.2 µm-filtered river water, photographed, and counted. Individual plastic pieces (3 - 50 mm) were gently rinsed and subdivided; one subsample was preserved in lysis buffer at −20 °C for DNA extraction. Water was sampled at all 14 stations; plastic debris was recovered by manta-net trawl at 13 of these, all except station GP2019-12 that was sampled for water only. Recovered plastic was not subdivided for DNA extraction at stations 02 and 04; plastisphere community and genome-resolved analyses are therefore based on the remaining 11 stations, with up to three plastisphere replicates preserved per station.

### Polymer characterization by ATR-FTIR and contamination control

Microplastics ≥ 300 µm were processed at the University of Basel under laminar-flow conditions. Samples were density-separated with NaBr solution, subjected to Fenton oxidation (FeSO₄ + H₂O₂) to remove organic matter, and vacuum-filtered onto a 300 µm PTFE membrane [5, 7, 8]. Putative microplastics were manually sorted under a stereomicroscope, mounted on double-sided adhesive tape (Tesa), and chemically characterized by ATR-FTIR spectroscopy using a Lumos microscope (Bruker Optics GmbH, Billerica, MA, USA) equipped with an MCT detector and germanium crystal (400–3350 cm⁻¹, 64 scans). Spectra were processed in R using OpenSpecy [25], applying polynomial baseline correction (degree 8) and Savitzky–Golay smoothing (polynomial 4, window 11). Particles were confirmed as plastic when baseline-corrected spectra produced concordant polymer assignments at match value > 0.85 (n = 7 stations with full ATR-FTIR characterization).

Two distinct microplastic metrics are reported: (i) total manta-net particle counts (n = 13 stations, unverified by polymer analysis); (ii) ATR-FTIR-confirmed microplastics ≥300 µm for a subsample of stations (n = 7 stations).

In the laboratory, a cotton lab coat was always worn, work was conducted under a laminar flow bench, and procedural blanks were run periodically throughout sample processing. PTFE wash bottles were used for transferring and cleaning samples (for detail see [7] Erni-Cassola et al., 2024).

### DNA extraction, library preparation, and sequencing

Biofilm material was extracted from lysis-buffer-preserved plastic particles using the Gentra Puregene Tissue Kit (Qiagen, Germany), following the manufacturer’s protocol as previously described [26]. DNA was quantified using the Quant-iT PicoGreen dsDNA Assay kit (Invitrogen, USA). Libraries were prepared with the Ovation Ultralow System V2 (Tecan, USA), size-selected for ∼350 bp inserts using AMPure XP beads and sequenced on an Illumina NextSeq 2000 platform at the Gulbenkian Institute of Science Genomics Unit (Oeiras, Portugal).

### 16S and 18S rRNA gene amplicon sequencing

The same DNA extracts used for shotgun metagenomic sequencing were also used for 16S rRNA gene (bacterial/archaeal) and 18S rRNA gene (eukaryotic) amplicon sequencing as previously described [27]. Briefly, amplicons were generated in technical triplicate using barcoded primers targeting the 16S rRNA gene (515F/926R, three-domain primer set, [28, 29]) and the 18S rRNA gene (515F/951R, [30, 31]). Triplicate PCR reactions per sample were pooled in equal volumes and purified with Agencourt AMPure XP beads (Beckman Coulter) prior to library pooling, and sequenced on the same Illumina NextSeq 2000 platform described above.

Read quality was assessed with FastQC, and paired-end reads were merged with PEAR (minimum overlap 10 bp). Merged reads were demultiplexed and primers removed via Cascabel v3.0 [32], with quality trimming, filtering, and amplicon sequence variant (ASV) inference performed using DADA2. Bacterial and archaeal taxonomy was assigned using SILVA v138.1 [33], and eukaryotic taxonomy using PR2 v5 [34]. 16S amplicon sequencing yielded usable taxonomic data for 12 of 14 water stations and 18S amplicon sequencing for 13 of 14 water stations; station-specific exceptions are detailed in the Results. ASV tables were built after removing singleton reads. Negative PCR controls and mock community standards run alongside the biological samples were used to decontaminate the resulting ASV tables with microDecon [35], with chloroplast- and chlorophyll-related ASVs additionally removed from the 16S dataset.

### Read quality control and assembly

Raw reads were processed with Cutadapt v3.1 [36] and fastp v0.20.1[37] to remove adapters, low-quality bases (Phred < 20), and low-complexity regions. Quality was assessed with FastQC v0.11.9 [38]. Cleaned reads were assembled de novo with MEGAHIT v1.2.9 [39] (default parameters; minimum contig length 1,000 bp). Quality assembly was evaluated with QUAST v4.6.3 [40].

### Metagenome-assembled genome reconstruction and classification

Metagenome-assembled genomes were provided by the Amaral-Zettler group at NIOZ, generated by independent binning of assembled contigs using MetaBAT2 v2.12.1 [41], MaxBin2 v2.2.4 [42], and CONCOCT v1.1.0 [43], with bin sets integrated and dereplicated with DAS Tool v1.1.0 [44]. Genome completeness and contamination were assessed with CheckM v1.2.4 [45], applying a stringent quality threshold of ≥70% completeness and ≤10% contamination, exceeding the standard MIMAG medium-quality criterion of ≥50% completeness [46]. Taxonomic assignment used GTDB-Tk (Galaxy wrapper v2.6.1+galaxy0) against the GTDB reference database release 226 (downloaded 2026-02-25) [47]. Reads were mapped to all 302 originally recovered MAGs using Bowtie 2 v2.5.1 [48]; resulting SAM files were sorted and indexed with samtools, and per-MAG coverage was aggregated using a custom Python script. Relative abundance was normalized across the full set of 302 MAGs before subsetting to genomes passing the quality threshold above, preserving each MAG’s true share of the whole recovered community; downstream analyses were restricted to the resulting 120 quality-filtered MAGs.

### Functional annotation

Protein-coding genes were predicted and annotated with DRAM v1.4.5 [49], drawing on KEGG [50], COG [51], and Pfam[52] databases (gene prediction via Prodigal v2.6.3 [53], internally by DRAM). Station-level functional profiles were computed by multiplying the MAG relative abundance matrix (MAG × station) by a KO presence–absence matrix (MAG × KO), yielding an abundance-weighted station × KO functional potential matrix (649 KOs × 120 MAGs).

### Statistical analyses

Environmental variables (PO₄³⁻, NO₃⁻+NO₂⁻, Si) were harmonized across all 14 stations ; NO₂⁻ alone was excluded to avoid redundancy with total oxidized nitrogen. All continuous variables were log₁₀-transformed and z-standardized prior to multivariate analyses.

Plastic abundance was standardized to volumetric concentration (particles m⁻³); and analyzed using two metrics separately: total manta-net particles (n = 13 stations) and ATR-FTIR-confirmed microplastics ≥300 µm (n = 7 stations). Station GP2019-12 was excluded from plastic analyses owing to missing volume data but retained for nutrient analyses. Associations between plastic metrics and dissolved nutrients were evaluated using Kendall’s rank correlation (τ), appropriate given the small sample size and tied ranks.

Community structure was assessed from Bray–Curtis dissimilarities of Hellinger-transformed ASV profiles and ordinated by NMDS (metaMDS(), vegan v2.6-4 [54]; set.seed(999); stress = 0.077) across all 23 station-habitat combinations (14 stations, with some stations contributing both plastisphere and water samples and others contributing only one). Environmental vectors (dissolved nutrients and total plastic concentration) were fitted to this ordination using envfit() (vegan v2.6-4; 9,999 permutations), applied separately to the plastisphere (n = 11) and water (n = 9) subsets, allowing direct comparison of nutrient– community coupling strength between habitats. Minimum sample-size requirements for envfit (n > 4k+1 for k ordination dimensions) precluded a separate envfit analysis restricted to ATR-FTIR-confirmed microplastic stations alone (n = 7); this metric was assessed by Kendall’s τ only (above). Bootstrap resampling (5,000 iterations, stations resampled with replacement) assessed the consistency of habitat differences in envfit R².

Alpha diversity was estimated for each station-habitat sample as Shannon diversity and observed ASV richness (diversity() and specnumber(), vegan v2.6-4), calculated from station-level count matrices after summing technical replicates; differences between habitats were tested using two-sided Wilcoxon rank-sum tests. Differences in taxonomic community composition between habitats were tested by PERMANOVA (adonis2(), vegan v2.6-4; 9,999 permutations) on Bray–Curtis dissimilarities of Hellinger-transformed ASV profiles from the 11 stations where both habitats were sampled (22 samples), with multivariate dispersion assessed using betadisper() (9,999 permutations).

Functional composition was calculated as the relative-abundance-weighted sum of DRAM-annotated KO profiles across the 120 quality-filtered MAGs in each station-habitat sample. Differences in functional composition between habitats were tested by PERMANOVA (adonis2(), 9,999 permutations), with permutations constrained by station (strata argument) to account for pairing. Multivariate dispersion was assessed using betadisper() followed by permutest() (9,999 permutations). Individual functional categories were screened for habitat-specific dispersion differences using the same approach, with FDR correction (Benjamini–Hochberg) across all categories. The aerobic corrin ring synthesis pathway was tested directly for a habitat difference in total abundance using a two-sided Wilcoxon rank-sum exact test.

All analyses were performed in R v4.5.2.

## Results & Discussion

### Microplastic abundance and polymer composition along an urban Rhine transect

Plastic debris was recovered at all 13 manta-net stations spanning the Rhine main stem. Total plastic concentrations co-varied significantly with dissolved oxidized nitrogen (NO₃⁻+NO₂⁻; Kendall τ = 0.587, p = 0.012) and silicate (τ = 0.624, p = 0.008), consistent with both plastic debris and nutrient enrichment tracking the same land-use gradient along the river continuum, though this association is correlative and does not establish a shared causal driver (Supplementary Table S2). Phosphate showed no significant association with total plastic abundance (τ = 0.367, p = 0.118). However, ATR-FTIR-confirmed microplastics (≥ 300 µm) at the seven characterized stations showed no correlation with any of the three nutrients (τ ≈ 0.048, p = 1.000 for all comparisons; Supplementary Table S2). This reflects the fact that verified particle counts were highest at mid-transect stations rather than declining consistently from upstream to downstream, in other words, the spatial pattern of ATR-confirmed particles did not follow the river gradient in the same way total plastic counts did, suggesting a genuinely patchy distribution rather than a sample-size limitation, though the smaller subsample (n = 7) warrants some caution in interpretation. Although there are similar reports of microplastics and total plastics showing diverse, patchy, non-linear distribution patterns along river transects, linked to proximity to urban areas rather than an upstream-downstream decline [55, 56], our subsample size might be too small to draw significant conclusions. Polymer characterization showed that PP, PE, and PS together accounted for the majority of identified particles (Fig. 1C), consistent with dominant polymer types reported in prior Rhine monitoring surveys [5, 7, 8] and across other major European rivers [22, 57]. This composition reflects the broad polymer mix expected from scattered urban and industrial sources distributed throughout the catchment. Together, these results establish the spatial and chemical context for the biological analyses that follow.

**Figure 1.**
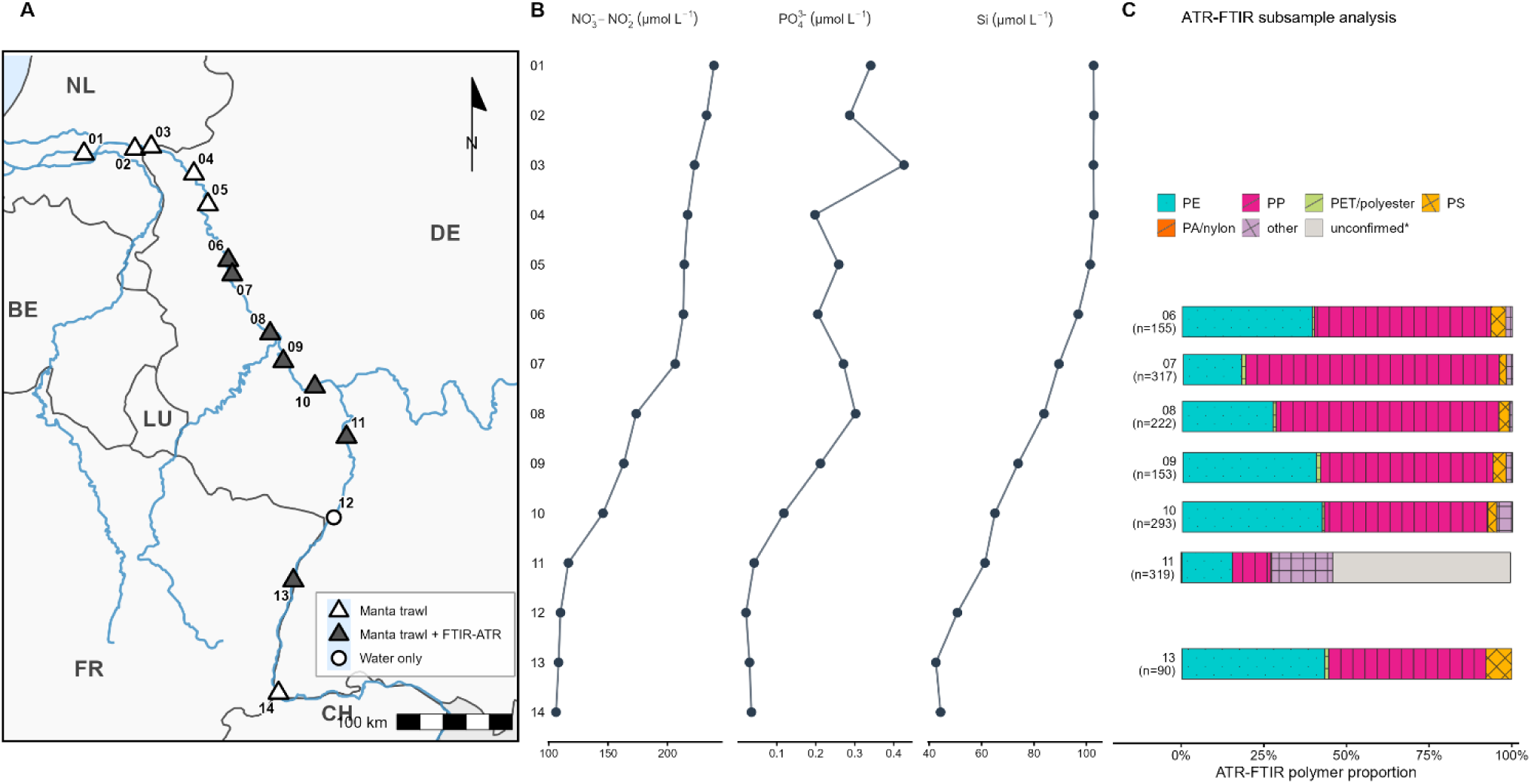
A) Location of 14 sampling stations spanning 820 km of the River Rhine from Rotterdam, NL (01) to Basel, CH (14). Symbol type indicates sampling method: open triangles, manta trawl only; filled triangles, manta trawl with ATR-FTIR polymer characterization; open circles, water sampling only. B) Dissolved oxidized nitrogen (NO₃⁻+NO₂⁻), phosphate (PO_4_^₃⁻^), and silicate (Si) concentrations across all 14 stations. C) Polymer composition of ATR-FTIR-confirmed microplastics at a subsample of seven stations for ATR-FTIR characterization, expressed as proportion of total characterized particles (n per station indicated). Colors and patterns indicate polymer type: PE, polyethylene; PP, polypropylene; PET/polyester, polyethylene terephthalate; PS, polystyrene; PA/nylon, polyamide; other, confirmed synthetic polymer not assignable to major categories; unconfirmed*, particles visually consistent with the other major microplastic particles at station 11 but not subjected to ATR-FTIR analysis.

### Plastisphere and water communities track the Rhine nutrient gradient with differing sensitivity

NMDS of ASV-level Bray–Curtis dissimilarities revealed a pronounced longitudinal gradient in community structure across all 23 station-habitat combinations (stress = 0.077; Fig. 2A). Dissolved oxidised nitrogen (NO₃⁻+NO₂⁻) was the strongest predictor of plastisphere community composition (envfit R² = 0.64, p = 0.012), with silicate and phosphate showing moderate but only marginally significant associations (R² = 0.48 and 0.47, respectively; p ≈ 0.07; Supplementary Table S2). Similarly, total plastic concentration showed a moderate, borderline significant association with the plastisphere ordination (R² = 0.51, p = 0.053; Supplementary Table S2). ATR-FTIR-confirmed microplastic counts were performed at only seven stations, precluding reliable envfit analysis given minimum sample size requirements for two-dimensional ordination (n > 4k+1 = 9; Kruskal [54]), and were therefore assessed by Kendall τ correlation only (Supplementary Table S2).

**Figure 2.**
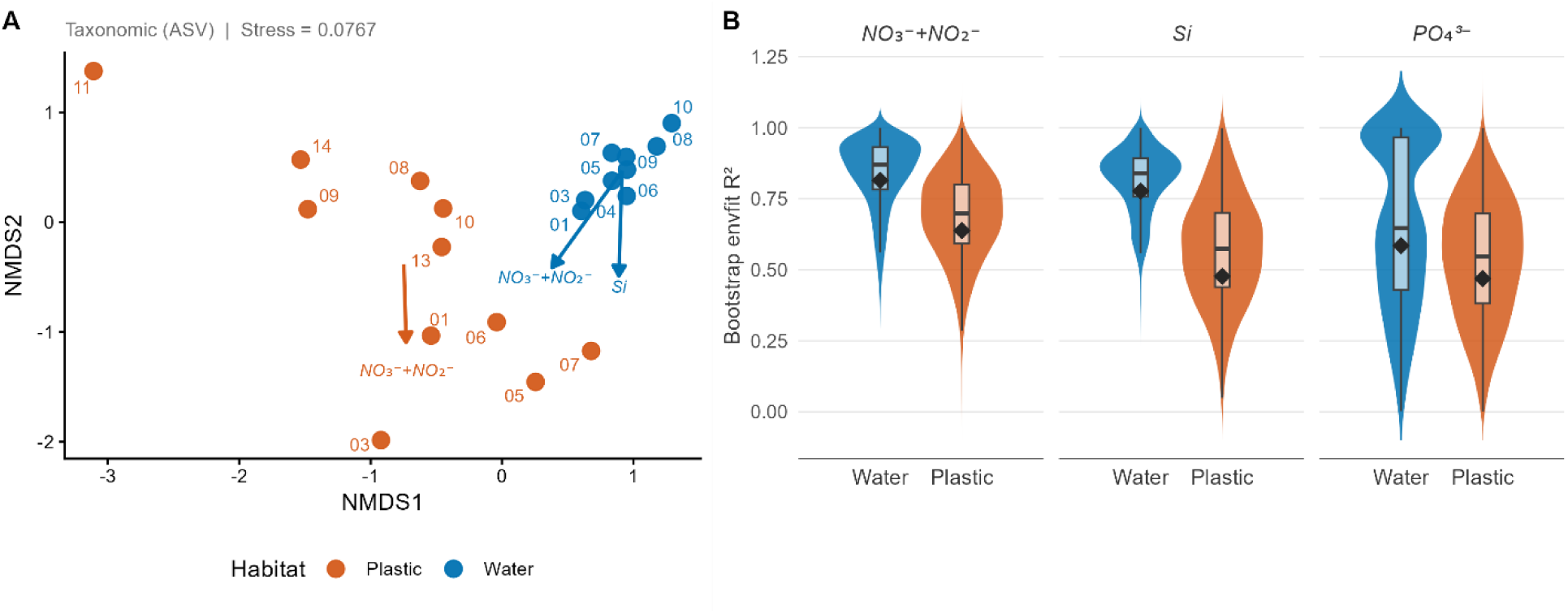
Plastisphere and water communities track the Rhine nutrient gradient with differing sensitivity. A) NMDS ordination of ASV community composition (Bray–Curtis dissimilarity, Hellinger-transformed; stress = 0.077) across 23 station-habitat combinations. Points represent individual station-habitat samples; colours indicate habitat (Water, blue; Plastic, orange). Environmental vectors show significant nutrient associations fitted by envfit (9,999 permutations; p < 0.05 shown only), run separately for plastisphere (MT, n = 11 stations; orange arrow) and water (ST, n = 9 stations; blue arrows) subsets; arrow length is proportional to R². B) Bootstrap distributions (5,000 resamples) of envfit R² for NO₃⁻+NO₂⁻ and Si, comparing coupling strength between water and plastisphere communities. Boxes show interquartile range and median; diamonds show observed R² from the full dataset.

Water communities showed markedly stronger alignment with the same dissolved nutrient gradients (NO₃⁻+NO₂⁻: R² = 0.82, p = 0.004; Si: R² = 0.78, p = 0.005; Supplementary Table S2). Bootstrap resampling (5,000 iterations; Fig. 2B) confirmed that this difference was consistent; water communities showed higher NO₃⁻+NO₂⁻–community R² than plastisphere communities in 80% of resample, and higher Si–community R² in 87% of resamples. Although 95% bootstrap confidence intervals overlapped zero, the direction and consistency of these differences support the interpretation that water communities are more tightly coupled to dissolved nutrient variation than their plastic-associated counterparts.

The weaker but still significant correlation of plastisphere communities to the dissolved nutrient gradient is consistent with evidence from large river transects that particle-associated and free-living microbial fractions respond differently to environmental chemistry [58]. For instance, along the Mississippi River, particle-associated communities responded to the nutrient gradient less tightly than their free-living counterparts, with dissolved inorganic nitrogen explaining substantially more variation in free-living than particle-associated communities [58], a pattern similar to our Rhine data. Two non-mutually-exclusive mechanisms may explain this difference: first, the plastic surface itself creates a physically and chemically distinct microhabitat [12, 15, 59]. It has been demonstrated that estuarine surface-associated biofilms generate steep oxygen gradients and accumulate reactive nitrogen intermediates, selecting for functional communities, including cobalamin-synthesizing comammox nitrifiers, that are metabolically decoupled from the surrounding water column [59]. Second, there is evidence that the order in which nutrient inputs reach a river community shapes its structure more persistently than the identity of the current nutrient load, because early-arriving populations gain competitive advantage through priority effects and microbial interactions mediated by amino acid and vitamin exchange [60]. Plastic surfaces, as stable floating substrates that retain biofilm communities across the stream gradient, may therefore accumulate community signal from multiple upstream conditions rather than responding exclusively to local dissolved nutrient concentrations. In this study, dissolved nutrient concentrations varied substantially across the transect, with NO₃⁻+NO₂⁻ ranging 2.3-fold (106.1–239.2 µmol L⁻¹), silicate ranging 2.4-fold (42.6–102.9 µmol L⁻¹), and phosphate ranging approximately 20-fold (0.021–0.426 µmol L⁻¹) across the 14 stations (Fig. 1 B), confirming a genuine longitudinal gradient capable of generating station-specific community signal for a stable, upstream-retaining substrate to accumulate.

Additionally, it has been found that stochastic processes play a larger role in structuring attached than free-living bacterial communities along dissolved nutrient gradients, including silicon [61], consistent with a biofilm environment that partially protects community assembly from the environmental chemistry. Together, these findings suggest that the partial decoupling we observe reflects both the physicochemical properties of the plastic surface (buoyant synthetic polymer) as a microhabitat and the legacy of upstream community assembly, neither of which is accessible to water communities that are continuously exposed to the ambient water column.

### 16S rRNA gene amplicon profiles confirm plastisphere taxonomic distinctiveness from planktonic communities

ASV-level community profiles derived from 16S rRNA gene amplicon sequencing confirmed that plastisphere and water communities are taxonomically distinct across all 11 stations where both habitats were sampled (22 samples total). PERMANOVA on Bray–Curtis dissimilarities showed that habitat explained 26% of community variation (R² = 0.258, F = 6.95, p = 0.0001). Alpha diversity did not differ significantly between habitats (water: Shannon diversity = 5.02 ± 0.48, observed ASV richness = 5,217 ± 1,943; plastic: Shannon diversity = 5.89 ± 1.11, observed ASV richness = 4,833 ± 4,628; Wilcoxon: Shannon W = 31, p = 0.056; Richness W = 79, p = 0.24), consistent with the unresolved debate in the literature over whether freshwater plastisphere communities harbor higher or lower diversity than their surrounding planktonic counterparts [62]. Plastisphere communities were, however, more variable in composition than water communities, with distances to group centroid approximately 1.5-fold higher on plastic surfaces than in the water column (betadisper: F = 79.57, p < 0.0001), suggesting that while the plastic surface selects for a broadly consistent set of taxa, local variation in colonization history, and surface conditioning (e.g., photodegradation) introduces a higher compositional heterogeneity than in the planktonic fraction [62, 63].

The two habitats differed markedly in dominant phyla. Water communities were dominated by Bacteroidota (40%) and Proteobacteria (30%), with Actinobacteriota (14%) and Cyanobacteria (10%) as secondary constituents, a composition consistent with that reported for free-living planktonic communities in large European rivers [22, 64]. Plastisphere communities were overwhelmingly dominated by Proteobacteria (62%), with Bacteroidota (15%), Planctomycetota (3%), and Acidobacteriota (3%) as minor components. The shift from Bacteroidota and Actinobacteriota-rich planktonic communities to a Proteobacteria-dominated biofilm mirrors patterns reported across freshwater plastisphere studies in rivers and headwater streams [64, 65], where Proteobacteria consistently dominate plastic-associated biofilms while Actinobacteriota and Bacteroidota remain more abundant in the surrounding water, and is consistent with the clear niche partitioning between plastisphere and surrounding water communities reported across nine major European rivers [22]. The enrichment of Proteobacteria on plastic surfaces reflects their metabolic versatility and capacity for surface attachment [16, 66, 67].

At a finer taxonomic resolution, water communities were dominated by a small number of genera: *Flavobacterium* (17.4%), *Limnohabitans* (11.3%), and *Pseudarcicella* (9.5%) together accounted for over a third of classified reads. By contrast, no single genus exceeded 6.5% relative abundance on plastic surfaces, despite Proteobacteria’s strong phylum-level dominance there. The most abundant plastisphere taxa comprised an unclassified genus within Comamonadaceae (6.5%), *Sphingorhabdus* (4.9%), *Flavobacterium* (4.4%), *Pseudomonas* (4.2%), and *Methylotenera* (4.0%), with several additional Comamonadaceae genera (*Rhodoferax*, *Aquabacterium*, and *Hydrogenophaga*) each contributing a further 2–3%. This varied genus composition suggests that surface attachment on plastic favors a taxonomically broad range of lineages within Proteobacteria, rather than a small number of taxa as seen in the water column.

Within the microeukaryotic plastisphere (Supplementary Table S3), pennate diatoms (Bacillariophyceae, mainly *Navicula* and *Cocconeis*) reached a higher relative abundance on plastic than in water (23.2% vs. 1.4% of eukaryotic reads), although this difference did not reach significance in a paired comparison across matched stations (Wilcoxon signed-rank, p = 0.11, n = 8), while centric, planktonic diatoms (dominated by *Stephanodiscus*) instead dominated the water column, reflecting the contrasting ecology of surface-attached and free-floating diatom lineages [15]. Sessile peritrich ciliates (e.g., *Epistylis*, *Zoothamnium*, *Vorticella*) were significantly enriched on plastic (Mann-Whitney, p = 0.02), further supporting this community as a surface-attachment specialist assemblage [65, 68]. Fungi (Ascomycota and Basidiomycota, dominated by *Trechispora*) were also strongly and significantly enriched on plastic (15.3% vs. 0.2% of eukaryotic reads; Mann-Whitney, p < 0.0001), consistent with fungal colonization of an already-established biofilm surface [69–71]. This enrichment considerably exceeds the fungal contribution reported for floating plastics across nine European rivers in a recent pan-European survey [72], where fungi remained a minor component of the microeukaryotic community (<5% of reads) and dominant genera did not include *Trechispora*, suggesting the Rhine plastisphere may host a locally distinct or more strongly plastic-associated fungal composition. Eukaryotic ASV richness and Shannon diversity (Supplementary Table S4, Figure S1) were both markedly lower on plastic than in water (Wilcoxon, p < 0.0001 for each, robust to rarefaction), suggesting that plastic substrates select for a narrower subset of colonizing taxa. The zebra mussel *Dreissena rostriformis* showed a significantly different distribution between habitats (Mann-Whitney, p = 0.0055), driven by a pronounced elevation on plastic at station 13 (Strasbourg), the site of a major inland Rhine port complex, and to a lesser extent at station 07 (Wesseling); at stations where paired water samples were available (07, 08), *Dreissena* signal was higher in water than on plastic; amplicon-based detection cannot distinguish whether this reflects planktonic larvae, shed eDNA, or both, rather than substrate-specific colonization [55].

Water samples at Stations 13 and 14 yielded negligible reads (0 and 1, respectively) in the eukaryotic (18S) amplicon dataset, despite adequate depth in the equivalent bacterial/archaeal (16S) amplicon dataset for the same samples; no eukaryotic comparison was possible at these two stations. The bacterial/archaeal (16S) amplicon dataset did not yield usable data for the Station GP2019-02 water sample, despite this sample succeeding in the equivalent eukaryotic (18S) amplicon dataset; Station 2 is therefore absent from the 16S taxonomic dataset. The Station GP2019-01 water sample was not included in the16S amplicon library preparation and sequencing, and is therefore also absent from the 16S taxonomic dataset, independent of the Station 2 exclusion described above. Stations 1 and 2 nonetheless remain part of the nutrient and total plastic abundance analyses reported elsewhere (see Methods), from which they were not excluded.

As diatoms cannot synthesize vitamin B_12_ (Cobalamin) and instead depend on co-occurring bacteria to meet this requirement [15, 19, 73, 74], the persistence of a diatom-rich, low-diversity eukaryotic plastisphere community raises the possibility that corrinoid provisioning by plastisphere bacteria is a key factor shaping this assemblage. To explore this relationship further, we conducted an exploratory Spearman correlation network between diatom genus relative abundances and aerobic corrin ring synthesis pathway abundance across the seven paired plastisphere stations (Supplementary Figure S2). Given the small sample size (n = 7), this analysis is intended as descriptive rather than confirmatory; nonetheless, all diatom genera examined showed positive correlations with aerobic corrin ring synthesis, consistent with a shared association between diatom presence and B_12_-related bacterial function on plastic surfaces.

### Functional potential on plastic is heterogeneous and converges on corrinoid biosynthesis

To characterize how plastic substrates reshape the metabolic potential of the plastisphere, we constructed a community-weighted functional profile from 120 quality-filtered MAGs and compared plastic-associated and water communities across the seven stations with paired MAG coverage data. At the level of the overall functional profile, habitat did not produce a significant shift in community centroid (PERMANOVA, R² = 0.245, F = 3.90, p = 0.125), but plastic-associated communities were significantly more variable in functional composition across stations than water communities (permutest, p = 0.0047), with mean distance to centroid roughly three-fold higher on plastic than in water (0.083 vs. 0.020 at the functional-category level). This indicates that plastic substrates do not impose a single, uniform functional signature across the transect, but instead generate substantially more site-to-site variability in metabolic potential than is seen among free-living communities.

Screening individual functional categories for habitat-specific dispersion differences (FDR-corrected across all categories) identified eight categories with significantly greater heterogeneity on plastic than in water (Fig. 3), most prominently CRISPR-associated defense systems and short-chain fatty acid/alcohol conversions (both p_fdr < 0.001), alongside three functionally related categories directly tied to vitamin B_12_ metabolism: aerobic corrin ring synthesis, anaerobic corrin ring synthesis, and adenosylcobalamin (ADO-CBL) biosynthesis (p_fdr = 0.010–0.034; Table 1). Among these, aerobic corrin ring synthesis, the core biosynthetic route for vitamin B_12_ (cobalamin) [20, 21], stood out for showing not only elevated dispersion but also a consistent, direction-specific enrichment: total pathway abundance was higher on plastic than in water at every one of the seven paired stations (Wilcoxon exact test, W = 3, p = 0.004079), with dispersion in this pathway roughly six-fold higher on plastic than in water (mean Bray–Curtis distance to centroid: 0.101 vs. 0.017; p_fdr = 0.010). Unlike the defense, and metabolism-related categories that showed heterogeneity without a clear directional pattern, corrinoid biosynthesis potential was both consistently elevated and more variable on plastic, indicating genuine restructuring of this specific metabolic function.

**Figure 3.**
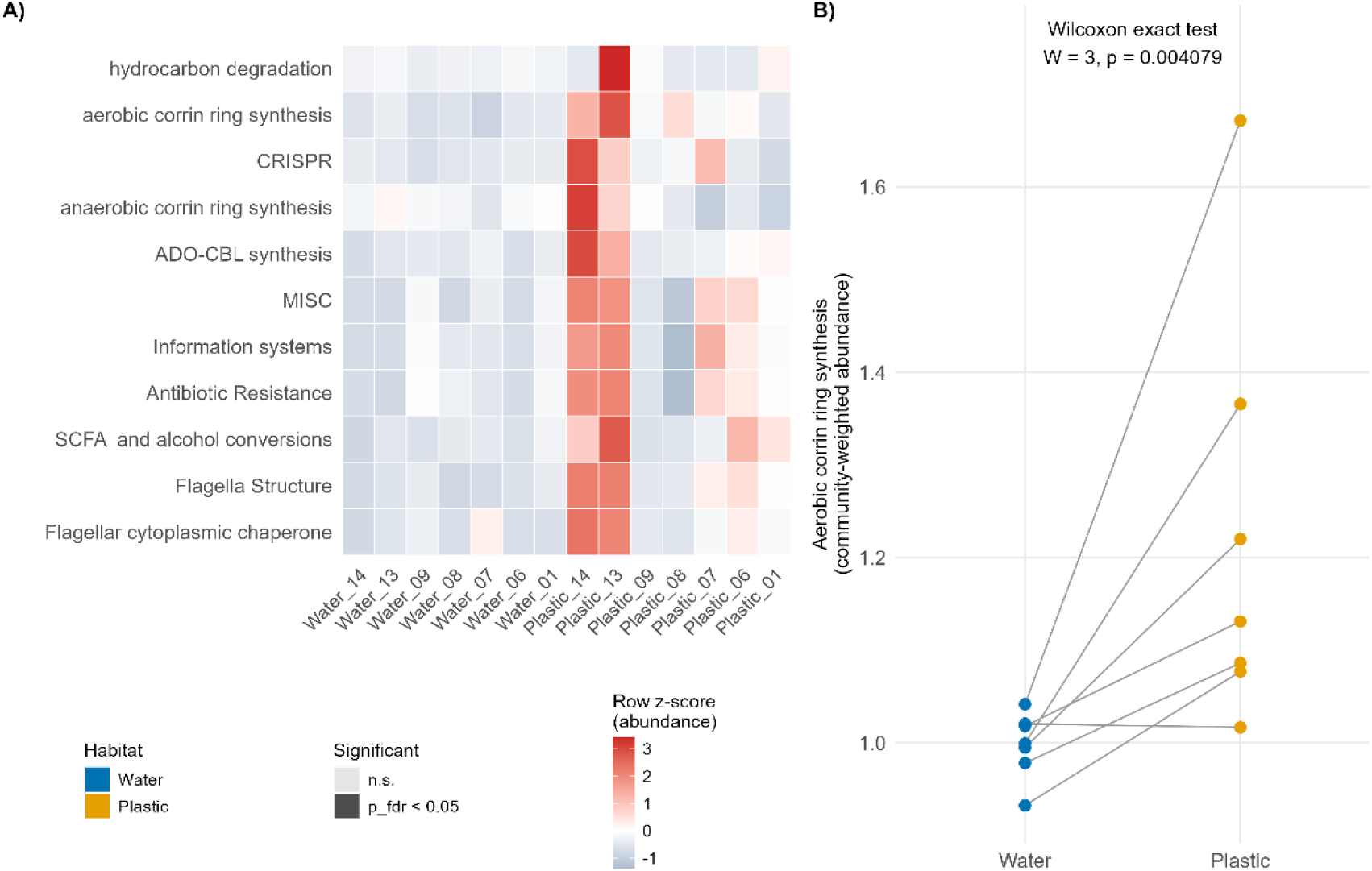
Functional potential on plastic is heterogeneous and converges on corrinoid biosynthesis. A) Community-weighted abundance of DRAM-annotated functional categories across 120 quality-filtered MAGs, 14 samples (7 paired stations). Rows ranked by Plastic:Water dispersion ratio; left strip marks FDR-significant categories (p_fdr < 0.05). B) Aerobic corrin ring synthesis (vitamin B12 biosynthesis) abundance, paired by station; higher on plastic at all 7 stations (Wilcoxon, W = 3, p = 0.004079).

**Table 1.** PERMANOVA and betadisper results for genome-resolved functional potential across plastisphere and free-living water communities.

| Test | R<sup>2</sup> | F | p-value |
| --- | --- | --- | --- |
| PERMANOVA: Habitat (plastic vs water) | 0.245 | 3.90 | 0.125 |
| betadisper: Plastic dispersion (mean BC) | ~0.090 | — | — |
| betadisper: Water dispersion (mean BC) | ~0.030 | — | — |
| permutest (dispersion difference) | — | — | 0.0047** |
| PERMANOVA: Station identity | 0.063 | — | 0.396 |
| PERMANOVA: NO<sub>3</sub><sup>-</sup>+NO<sub>2</sub><sup>-</sup> | 0.073 | — | 0.355 |
| PERMANOVA: PO<sub>4</sub><sup>3-</sup> | 0.111 | — | 0.253 |
| PERMANOVA: Si | 0.034 | — | 0.579 |

Significant results indicated by \*\* $p < 0.01$ . BC= Bray-Curtis

This finding is consistent with the mechanistic framework proposed for marine plastisphere biofilms, in which cobalt and zinc transporters and methylmalonyl-CoA mutase, enzymes central to B_12_-dependent metabolism, become more abundant as plastisphere biofilms mature [15, 75]. Diatoms cannot synthesize vitamin B_12_ de novo and instead depend entirely on co-occurring bacteria to meet this requirement [15, 58–60], directly linking this result to the diatom-rich, low-diversity eukaryotic plastisphere community established above. The persistence of pennate diatoms on a plastic surface may therefore depend on the concurrent enrichment of corrinoid-producing bacteria, positioning plastic substrates as a habitat that actively provisions, rather than merely hosts, a metabolically interdependent community. Taken together, these results indicate that plastic surfaces restructure both the taxonomic composition and the specific metabolic capacities of the communities they support, rather than passively accumulating biomass from the surrounding river.

Together, these results indicate that plastic substrates in the Rhine impose two distinct but related consequences for microbial community assembly: a partial decoupling of taxonomic structure from the dissolved nutrient gradient that otherwise governs the surrounding water column, and a targeted restructuring of specific metabolic capacity, most notably, a consistent enrichment of corrinoid biosynthesis potential across all seven paired stations. Unlike the broader taxonomic and functional heterogeneity we observe, which varies in direction and magnitude across sites, the corrinoid signal is both directionally consistent and among the most differentially dispersed functions we detected, suggesting it reflects a genuine, substrate-associated selective pressure rather than incidental site-to-site noise. The patterns described here are properties of the plastic-biofilm habitat as a whole, its persistence as a discrete, surface-associated microenvironment distinct from the planktonic water column, rather than evidence that particular polymer types select for particular communities or functions.

Several caveats warrant consideration. This cross-sectional design captures spatial variation at a single time point; temporal replication would help establish whether the taxonomic and functional patterns described here persist across seasons and flow conditions or are specific to the sampling period. The ATR-FTIR-confirmed microplastic subset is limited to seven stations, reducing statistical power for polymer-specific comparisons; future campaigns would benefit from polymer characterization at every station. MAG-based functional profiles represent the assembled, higher-biomass fraction of the plastisphere, and coverage-based abundance estimates may under-represent rare or low-biomass colonizers relative to more sensitive amplicon-based approaches. Finally, other environmental variables such as temperature, turbidity, dissolved organic carbon, and surface weathering state were not fully characterized at every station and may contribute additional explanatory power in future, more heavily instrumented sampling designs.

In conclusion, this study provides genome-resolved evidence that plastic substrates in a large European river host taxonomically and functionally distinct microbial communities, with community assembly on plastic only partially explained by the dissolved nutrient gradients that structure the surrounding water column. Central to this restructuring is a consistent enrichment of vitamin B_12_ biosynthetic potential, linking the plastisphere’s altered functional capacity directly to the diatom-rich, low-diversity eukaryotic community it supports. Future work combining longitudinal sampling with controlled incubations would help disentangle the relative contributions of substrate persistence, biofilm maturation, and local environmental conditions to this restructuring, and transcriptomic or metabolomic follow-up would clarify whether the corrinoid biosynthesis potential identified here is constitutively expressed or induced by the plastisphere’s diatom-bacteria interactions. More broadly, understanding how plastic substrates reshape microbial metabolic capacity in rivers is a necessary step toward assessing the biogeochemical and ecological consequences of the plastic continuum linking terrestrial and marine ecosystems.

## Acknowledgements

We thank Greenpeace Netherlands and Greenpeace Germany for providing access to the research vessel Beluga II and for logistical support during the 2019 Rhine River expedition, which made sample collection across the 820 km transect from Rotterdam to Basel possible.

The authors thank Ms. Nicole Daniela Seiler-Kurth and Miss Sara Staubli from the Man-Sociaty-Environment Group at the University of Basel, as well as Mr. Nikolai Matviiets and Mr. Alejandro Abdala from NIOZ, Netherlands for their invaluable technical support, and Christine Gogel for her administrative support throughout the project.

This work was supported by a donation from the Wilsdorf Mettler Future Foundation to the University of Basel in support of the research initiative “Mikroplastik im Rhein.” We thank Dr. Jörg Kübel and Dr. Igor Korneitchouk of the Wilsdorf Mettler Future Foundation for their support. The funder had no role in the design, execution, analysis, or interpretation of the study, or in the decision to submit the article for publication.

## Author contributions

Priscilla Carrillo-Barragán (Methodology, Investigation, Formal Analysis, Writing—original draft, Writing—review & editing), Erik Zettler (Conceptualization, Methodology, Investigation, Writing— review & editing), Sebastian Rieder (Methodology, Investigation, Formal Analysis), Leslie G Murphy (Methodology, Investigation, Formal Analysis), Tom Theirlynck (Methodology, Investigation, Formal Analysis), Patricia Burkhardt-Holm (Conceptualization, Methodology, Writing—review & editing, Supervision, Funding Acquisition), and Linda Amaral-Zettler (Conceptualization, Methodology, Formal Analysis, Investigation, Writing—review & editing, Supervision, Funding Acquisition).

## Data availability

Raw sequence data generated for this study are publicly available in the National Centre for Biotechnology Information (NCBI) database under BioProject PRJNA1506697. Water and plastisphere samples used for community profiling are deposited as individual BioSamples under the same BioProject (14 water samples, accessions SAMN62142635–SAMN62143646; 11 plastisphere samples, accessions SAMN62145081– SAMN62145091). The metagenome-assembled genomes (MAGs) reconstructed and analyzed in this study (n = 120) are deposited as individual BioSamples under the same BioProject (accessions SAMN62216404– SAMN62216534), with corresponding genome assemblies deposited in GenBank. The codes used for analyses and figure generation are available in the GitHub repository at https://github.com/PrisCB/Beluga-Rhine-Plastisphere.

