## SupplementaryMaterials for "Riverine plastic litter restructures microbial vitamin B_12_ metabolism and generates functional heterogeneity"

**Supplementary Table S1.** Station-habitat combinations included in the 16S community ordination (n = 23)

| Combination | Original Station ID | Station | Habitat |
|---|---|---|---|
| 1 | GP2019-01 | 01 | Plastic |
| 2 | GP2019-03 | 03 | Plastic |
| 3 | GP2019-03 | 03 | Water |
| 4 | GP2019-04 | 04 | Water |
| 5 | GP2019-05 | 05 | Plastic |
| 6 | GP2019-05 | 05 | Water |
| 7 | GP2019-06 | 06 | Plastic |
| 8 | GP2019-06 | 06 | Water |
| 9 | GP2019-07 | 07 | Plastic |
| 10 | GP2019-07 | 07 | Water |
| 11 | GP2019-08 | 08 | Plastic |
| 12 | GP2019-08 | 08 | Water |
| 13 | GP2019-09 | 09 | Plastic |
| 14 | GP2019-09 | 09 | Water |
| 15 | GP2019-10 | 10 | Plastic |
| 16 | GP2019-10 | 10 | Water |
| 17 | GP2019-11 | 11 | Plastic |
| 18 | GP2019-11 | 11 | Water |
| 19 | GP2019-12 | 12 | Water |
| 20 | GP2019-13 | 13 | Plastic |
| 21 | GP2019-13 | 13 | Water |
| 22 | GP2019-14 | 14 | Plastic |
| 23 | GP2019-14 | 14 | Water |

Station-specific exclusions from the 16S taxonomic dataset (Stations 1, 2) are detailed in the Results. Station 4: plastic debris was recovered in the field but not subdivided for DNA extraction; no plastisphere sequencing library exists for either marker. Station 12: water-only station (no plastic collected); included here as one of the 12 water stations with usable 16S data.

**Supplementary Table S2.** Summary of envfit vector fitting and Kendall rank correlations between dissolved nutrients, total microplastic concentration, and community structure. envfit was performed using 9,999 permutations. ATR-FTIR-confirmed microplastic counts (ATR_MPs, n = 7) did not meet the minimum sample size requirement for envfit (n > 4k+1 for k = 2 ordination dimensions; Kruskal, as cited in Oksanen et al. 2022, vegan v2.6-4) and were assessed by Kendall τ only. TotalPlastic = Greenpeace total particle counts. ATR_MPs = ATR-FTIR verified particle counts at Universit of Basel *p < 0.05; **p < 0.01.

| Test | n | τ or $R^2$ | p-value | Habitat |
|---|---|---|---|---|
| ***envfit vector fitting — Plastisphere*** | | | | |
| envfit $NO_3^-$+$NO_2^-$ | 11 | 0.638 | **0.012*** | Plastisphere |
| envfit Si | 11 | 0.477 | 0.065 | Plastisphere |
| envfit $PO_4^{3-}$ | 11 | 0.469 | 0.067 | Plastisphere |
| envfit TotalPlastic | 11 | 0.511 | 0.053 | Plastisphere |
| ***envfit vector fitting — Water*** | | | | |
| envfit $NO_3^-$+$NO_2^-$ | 9 | 0.815 | **0.004**** | Water |
| envfit Si | 9 | 0.777 | **0.005**** | Water |
| envfit $PO_4^{3-}$ | 9 | 0.585 | 0.075 | Water |
| ***Kendall τ — TotalPlastic vs nutrients*** | | | | |
| TotalPlastic vs $NO_3^-$+$NO_2^-$ | 11 | 0.587 | **0.012*** | — |
| TotalPlastic vs $PO_4^{3-}$ | 11 | 0.367 | 0.118 | — |
| TotalPlastic vs Si | 11 | 0.624 | **0.008**** | — |
| ***Kendall τ — ATR_MPs vs nutrients (ATR-FTIR verified counts only)*** | | | | |
| ATR_MPs vs $NO_3^-$+$NO_2^-$ | 7 | 0.048 | 1.000 | — |
| ATR_MPs vs $PO_4^{3-}$ | 7 | 0.048 | 1.000 | — |
| ATR_MPs vs Si | 7 | 0.048 | 1.000 | — |

**Supplementary Table S3**. Eukaryotic (18S rRNA) community composition and diversity, Plastic vs. Water (Rhine transect).

Relative abundances are means across samples (n = 20 Plastic, n = 11 Water; one negative control and one failed sample [<100 total reads] excluded). Values are % of eukaryotic reads per sample, at Class level unless otherwise noted. Statistical tests are two-sided Mann-Whitney U (Wilcoxon rank-sum) tests on sample-level relative abundances. Where the pooled data contain no ties (Fungi; ASV richness; Shannon diversity), the exact permutation p-value is reported; all other tests use the asymptotic normal approximation with continuity correction, as ties preclude an exact calculation. Classes are listed by combined Plastic+Water mean abundance rather than by pre-selected taxa, so this table reflects the dominant community members overall, not only the groups discussed in the main text.

| Class / group | Plastic mean (%) | Plastic SD (%) | Water mean (%) | Water SD (%) | Test, p-value |
|---|---|---|---|---|---|
| Pennate diatoms (Bacillariophyceae) | 23.18 | 29.63 | 1.41 | 0.61 | p = 0.060 (unpaired); p = 0.11 (paired, matched stations n=8) |
| Sessile ciliates (Oligohymenophorea, incl. Peritrichia_2) | 20.62 | 23.51 | 1.79 | 0.61 | p = 0.030 (class-level); p = 0.022 (Peritrichia_2 order specifically) |
| Fungi (Ascomycota + Basidiomycota, dominated by Trechispora) | 15.26 | 23.28 | 0.19 | 0.17 | $p = 3.3\times10^{-6}$ (exact test) |
| Land-plant material (Embryophyceae) | 6.71 | 19.21 | 0.11 | 0.05 | p = 0.35 (not significant — driven by few outlier samples, SD>mean) |
| Annelida (worms) | 4.64 | 18.29 | 0.18 | 0.55 | p = 0.25 (not significant — same caveat) |
| Nematoda | 4.06 | 12.56 | 0.37 | 0.36 | p = 0.17 (not significant — same caveat) |
| Mollusca (incl. Dreissena rostriformis) | 2.35 | 5.86 | 3.72 | 3.76 | p = 0.017 |
| Other ciliates (Spirotrichea) | 2.33 | 6.57 | 8.97 | 5.85 | $p = 7.0\times10^{-5}$ (water-dominant) |
| Chrysophyceae (golden algae) | 0.89 | 1.31 | 15.61 | 10.80 | $p = 6.1\times10^{-6}$ (water-dominant) |
| Centric diatoms (Mediophyceae) | 0.04 | 0.12 | 32.98 | 11.57 | $p = 3.8\times10^{-6}$ (water-dominant) |
| Cryptophyceae | 0.01 | 0.04 | 6.01 | 2.39 | $p = 3.2\times10^{-6}$ (water-dominant) |
| Dreissena rostriformis specifically | 2.11 | 5.87 | 2.13 | 3.44 | p = 0.0055 (see distribution-shape note below) |
| Sum of groups shown above / all remaining classes combined | 80.1 / 19.9 | – | 71.3 / 28.7 | – | – |

Note on Embryophyceae, Annelida, and Nematoda: despite high mean relative abundance on plastic, none of these groups differ significantly between habitats. In each case the standard deviation exceeds or approaches the mean, indicating the signal is driven by a small number of samples with unusually high counts (likely incidental terrestrial detritus or metazoan bycatch) rather than a consistent colonisation pattern. These are reported for completeness but are not interpreted as enrichment.

Note on *Dreissena rostriformis:* on plastic, undetected in 9 of 20 samples but reaching up to 25.1% of reads elsewhere (patchy/bimodal distribution); in water, present at consistently low levels in all 11 samples (median 0.68%, never absent). Median relative abundance was therefore lower on plastic (0.03%) than in water (0.68%) despite similar means.

**Supplementary Table 4.** Eukaryotic Alpha diversity

| Metric | Plastic mean ± SD | Water mean ± SD | Test, p-value | Notes |
|---|---|---|---|---|
| ASV richness (raw) | 391 ± 358 | 2004 ± 318 | $p = 2.4\times10^{-8}$ (exact test) | n=20 Plastic, n=11 Water |
| Shannon diversity (raw) | 3.05 ± 0.99 | 4.90 ± 0.40 | $p = 2.4\times10^{-8}$ (exact test) | Complete separation (U=0) |
| ASV richness (rarefied to 36,071 reads) | 346 ± 292 | 1602 ± 189 | $p = 2.4\times10^{-8}$ (exact test) | 50 iterations/sample |
| Shannon diversity (rarefied) | 3.05 ± 0.99 | 4.88 ± 0.40 | $p = 2.4\times10^{-8}$ (exact test) | Confirms depth is not a confound |

Note: rarefaction to the minimum sample depth (36,071 reads; 50 resampling iterations per sample, seed = 42) reproduces the raw-count result almost exactly, confirming the diversity difference is not attributable to unequal sequencing depth (mean depth: Plastic 177,104 ± 52,808; Water 158,737 ± 54,445 reads; Mann-Whitney on depth itself, p = 0.24, not significant, asymptotic — ties present).

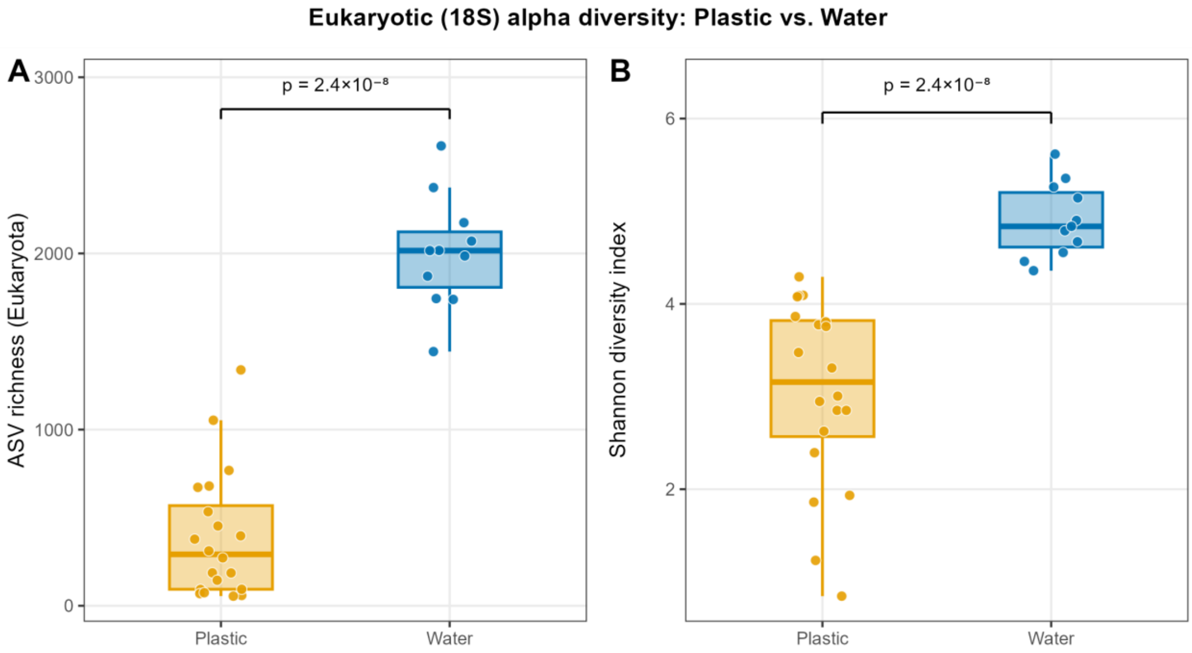


**Supplementary Figure S1**. Eukaryotic alpha diversity derived from 18S rRNA amplicons, comparing plastic and water samples across Rhine sampling stations.

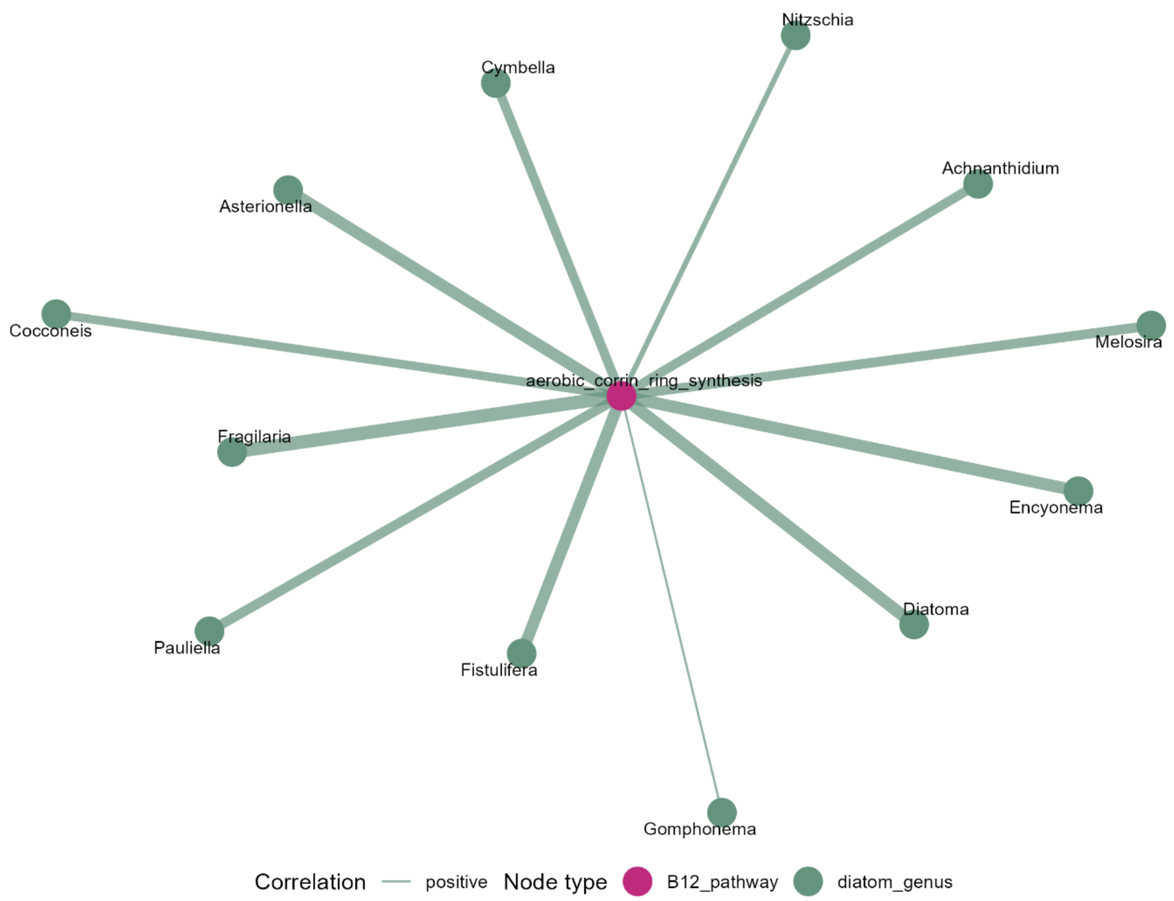


**Supplementary Figure S2**. Exploratory Spearman correlation network between diatom genus relative abundances and the aerobic corrin ring synthesis pathway across seven paired plastisphere stations (18S rRNA amplicon and metagenomic functional data, respectively). Edge width is proportional to correlation strength; edge colour indicates direction (positive, blue). Node colour distinguishes diatom genera (green) from the B12 biosynthesis pathway (pink).
